# In model-based fMRI significant is less than specific

**DOI:** 10.1101/429621

**Authors:** Erik J Peterson, Carol A Seger

**Affiliations:** Dept of Psychology, Carnegie Mellon University, Pittsburgh, PA; Psychology Dept, Colorado State University, Fort Collins, CO

## Abstract

By comparing computational model output to BOLD signal changes model-based fMRI has the potential to offer profound insight into what neural computations occur when. If this potential is to be fully realized, statistically significant outcomes must imply specific outcomes. That is, we must have a clear idea of how often a model not present in the BOLD signal but present in the predictor set will reach significance. We ran Monte Carlo simulations of reinforcement learning to examine this kind of specificity, focusing in on two aspects. One, to what degree can we tell related but theoretically distinct predictors apart. About 40% of the time the studied predictors were indistinguishable. Two, how well can we separate out different parameterizations of the same reinforcement learning terms. Nearly all parameter settings were indistinguishable. The lack of specificity between models and between parameters suggests a uncertain relation between significance and specificity. Follow up analyses suggest the temporally slow and prototyped nature of the haemodynamic response (HRF) can substantially increase correlations, ranging from −0.16 to 0.73 with an average of 0.27. Though we focused on a single case study, i.e., reinforcement learning, specificity concerns are potentially present in any design which does not account for the slow prototyped nature of the HRF. We suggest more specific conclusions can be reached by moving from null hypothesis testing approach to a model selection or model comparison framework.

## Introduction

The neural model implicit in many fMRI analyses is a simple switch. Regions of the brain turn on then off. As an example consider a simple learning experiment with two reward levels. To compare large rewards (e.g. “Win $10!”) to small rewards (e.g. “Win $0.01!”) one typically forms a “impulse”-based GLM design matrix with two columns. In the first column large rewards get coded as 1, while small rewards are coded as 0. In the second column small rewards have the opposite code, i.e., “Win $0.01!” gets coded as 1. Each impulse-based column is then convolved with a haemodynamic response function (HRF), regressed onto each voxel’s blood oxygen level dependent (BOLD) time course followed by a statistical contrast of the two reward conditions, along with a multiple comparison correction. This is the standard statistical parametric mapping (SPM) routine (Josephs et al., 1997) and it is relatively simple to understand and to implement. It is also robust to noise and other natural (e.g. regional) HRF shape variation (Henson et al., 2001; Friston et al., 1998), as thousands of reports empirically demonstrate (Bandettini, 2007).

In reality however the neural response and the resulting HRF is not an all or none function. The HRF changes in size and shape as function of stimulus, e.g. as a function of visual contrast (Boynton et al., 1996). In fact, stimulus and context dependent changes seem to be the norm. Reward valence and magnitude (Delgado et al., 2003, 2000), motivation (Delgado et al., 2004), response accuracy (Seger and Cincotta, 2005), strength of recall (Wais, 2008), degree of regret (Fujiwara et al., 2008), and many other tasks and conditions all show distinct variations in HRF shape.

Model-based fMRI tries to predict such shape variations by replacing impulse codes, where the HRF shape representing each trial’s response is identical, with varying trial-level estimates. We focus solely here on estimates derived from computational modeling efforts. These model-based designs are therefore specific hypotheses about *what* mathematical computation happened *when*. By comparing biologically plausible model implementations, model-based fMRI can be extend to examine *how* a computation occurred (O’Doherty et al., 2007; Mars et al., 2010). But to meaningfully estimate *what, when*, and *how* in an SPM framework, significant outcomes should imply specific outcomes. That is, there should be strong relationship between a *p* value crossing the significance threshold and the model data closely resembling the real data.

Predicting fMRI BOLD changes with modeling is implicitly a shift from qualitative methods, those focused on ‘neural signatures’ and broad task-related regional differences, to a quantitative method set capable of contributing to the crucial and ongoing theoretical discussions of regional neural computation. But for that shift to be fully realized, when a model-based predictor is significant that significance must also imply only numerically *very similar* alternative predictors would also be significant. Put another way, model-based designs must be able to reliably distinguish between theoretically distinct co-variates.

Impulse codes work well in practice because they act as a correlational catchall. In reality trial-by-trial HRFs vary, but most if not all those variations still correlate with the convolved impulse time course (Baumgartner et al., 2000). This is an extremely useful property for analyses focused on qualitative neural signatures of *where* activity occurred. But the success of impulse codes suggests a problem for model-based predictors. While model-based designs try to make specific trial-level magnitude predictions, the slow prototypical nature of the HRF may lead any given model-based predictor to mistakenly “catch” theoretically distinct signals, just like impulse designs. To better understand and quantify this potential specificity problem, we ran several Monte Carlo simulations of model-based fMRI.

We chose to focus on reinforcement learning models, making them a case study. Reinforcement learning measures are one of the most studied model-based predictors. Evidence for a conserved set of neural reinforcement learning signals comes from electrophysiological studies in multiple species (Mirenowicz and Schultz, 1994; Hollerman et al., 1998; Roesch et al., 2007), genetic variation studies (Frank et al., 2007) and causal modalities (Pessiglione et al., 2006; Pizzagalli et al., 2008; Frank and O’Reilly, 2006), and of course model-based fMRI studies (Glimcher, 2011; Montague et al., 2006; D’Ardenne et al., 2008; McClure et al., 2003; Seger et al., 2010; Garrison et al., 2013). This strength of evidence allows us to focus not on the truth of reinforcement learning theory *per se*, but instead on potential limits of model-based analyses using a well established body of work as our starting place.

## Material & Methods

We conducted a series of Monte Carlo simulations to examine the specificity of model-based fMRI. We employed Rescorla-Wagner reinforcement learning models, and other reward-related regressors, as a case study. First we will describe the construction of the simulated behavioral data. Second is reinforcement learning model construction and parameter fitting. Third we describe the mechanics of the fMRI simulations.

In the course of this and other work we developed a new fMRI simulation package for Python dubbed “modelmodel” (http://www.robotpuggle.com/code/). This library is unique both in its focus on model-based simulations and its ability to seamlessly intermix simulations with ROI analyses of real fMRI data. All other simulation code is available for download and (re)use. Each simulation consisted of 1000 iterations. Visualization and other post-simulation analysis was completed in R language.

### Simulating behavioral data

Behavioral data consisted of *N* = 60 simulated trials over a single condition *c* intermixed with 30 “baseline” jitter periods sampled from the the uniform distribution, *U* (1,6). Jitter periods are short random pauses that allow for rapid event-related fMRI designs, i.e. designs whose trials are shorter than the Ȉ 30 seconds it takes for the BOLD response to return to its baseline level.

We simulated two types of behavior – learning and guessing. During learning, behavior was modeled as guessing (i.e. *p ~ U* (0,1)) until some trial *k* after which learning began. Learning was simulated by sequential sampling of the cumulative normal distribution 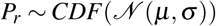. The mean (*µ*) was itself sampled from 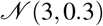 with the standard deviation (σ) fixed at 1. The transition point *k* shifted the onset of learning behavior, while *µ* modulated the slope of the learning curve. Parameter *k* was drawn from *U* (0,59), matching the (jitter-excluded) trial index (*i* = {0,1,2,···*N* – 1}). Sample guess was drawn only from the uniform distribution (i.e. *Pr ~ U* (0,1)). Accuracy data for trial *A* (*i*) was generated for both learn and guess using *P_r_* and the binomial distribution (e.g., *Alearn* (*i*) *~ B* (1*, plearn* (*i*)), Figure 1A and B). The parameters *k* and *µ* were adjusted until *P_r_* was broadly similar to learning curves we’ve observed in previous empirical studies of stimulus-response and reinforcement learning (Lopez-Paniagua and Seger, 2011; Seger et al., 2010; Seger and Cincotta, 2005) (Figure 1A). guess time courses had no notable dynamics; accuracy and *P_r_* fluctuated around 0.5 (Figure 1B). Trial counts ranging from 20 to 100 (*N*) were initially examined, but had no substantive impact on our results.

### Reinforcement learning models

Two approaches for generating reinforcement learning data were employed. The first, the *confusion* approach, assessed how likely different performance, reward probability, and reinforcement learning regressors might be confused in a GLM context. In the *confusion* analysis the learning rate *α* and response volatility parameter *β* were fit by maximum log-likelihood, as is typical in the behavioral literature (Glimcher, 2011; Montague et al., 2006; D’Ardenne et al., 2008; McClure et al., 2003; Seger et al., 2010).

For the *confusion* data, model parameters were exhaustively searched, in 0.01 increments. The learning rate *α* spanned (0,0.01,0.02*···*1). The choice parameter *β* spanned (0,0.01,0.02*,···*5). Log-likelihood estimates were calculated based on a softmax transform of trial-wise values *V_t_*(*c*) into the log of choice probabilities followed by a summation (Eq 3). The best parameter set was the one that maximized log-likelihood 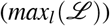. For example time courses see Figure 1A and B.

In the second approach, *α* values were selected *a priori*. This *separation* design assessed to what degree different parameterizations of the same reinforcement learning measures can be reliably separated in a model-based design. In the *separation* set *α* was iteratively set to 0.1,0.3,0.5,0.7,0.9 (see Figure 1B and C). As there was no search, no softmax transform was required and no *β* settings were considered. Only guessing behavioral models were employed for the separation analyses.

In both *confusion* and *separation* we examined two classic reinforcement learning measures. First was *value*, denoted *V_i_*(*c*), and calculated as in Eq 2. *Value* is an estimate of total future rewards, a measure closely tied to the expected value (Sutton and Barto, 1998). Second was the reward prediction error (denoted as *δ* or *RPE*, see Eq 1). *RPEs* result from the comparison of the current estimate of value to the received reward (Eq 1). The learning rate (*α*) controls how much the current *value* update is influenced by past updates. With lower *α* settings current learning is strongly affected by *past values*, leading to a slower progression. Large *α* values have the opposite effect; learning is fast but volatile and history has little effect (compare the leftmost and rightmost columns in Figure 1C). The salience parameter (denoted as *γ* here) was set to 1 for all analyses. 
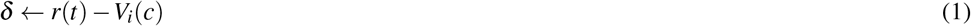

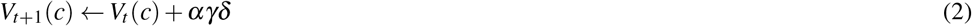

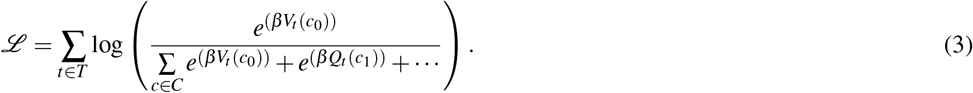

**Figure 1.**
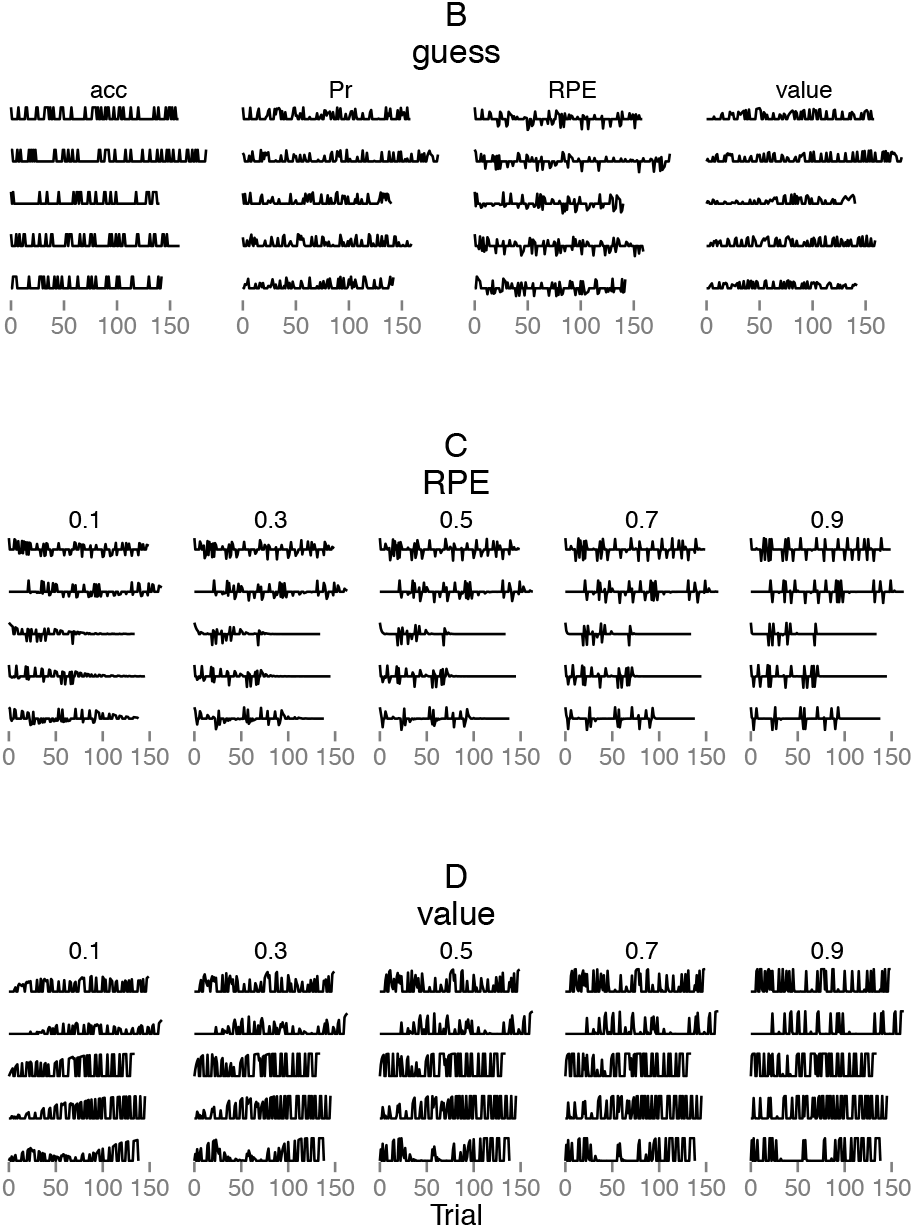
Randomly selected examples of the four simulated reinforcement learning measures – accuracy (acc), reward probability *P_r_*, an estimate of expected *value*, and reward prediction error *RPE*. **A** and **B** represent time courses for the *confusion* analysis and were determined, in part, based on the maximum likelihood fitting procedure described in the text. **A** is examples of behavioral learning. **B** shows simulated guessing behavior. Note how the *value* graphs show a general rise across trials as subjects learn, whereas *RPE* decreases in variability across learning as fewer errors are made. The bottom two panels are from the *separation* analysis and demonstrate how *RPE* (**C**) and *value* (**D**) change with learning rate (*α*). Each column in the bottom two panels matches a value of *α* ranging from 0.1 to 0.9 in 0.2 steps. In **C** and **D** all example data is from the learn behavioral models.

### fMRI simulations

FMRI data was constructed from behavioral and reinforcement learning time-series. Each series was convolved with the “canonical SPM” HRF. The canonical HRF is an impulse response characterized by two gamma functions, one for the peak and one for the post-peak undershoot. It is parameterized by a peak delay of 6 s, an undershoot of 16 s. The peak/undershoot amplitude ratio is 6 (Penny et al., 2006).

BOLD data was simulated by combining these HRF-convolved series with white noise 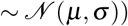. Several other noise sources were examined, including 1/*f*, autocorrelated white noise (AR(1)), white noise plus respiration confounds, and white-noise plus low frequency drifts. Each of these alternate noise sources reduced specificity more than white noise. In some cases (e.g. the low frequency drift and respiration models) the reduction was large, *>*25%. The *qualitative* pattern of results was, however, unaltered by noise. We went with the conservative choice of white noise. If specificity was low with white noise, the problem would only worsen with more realistic noise choices.

For the *confusion* analyses the model-based predictors were *value* and *RPE*. The behavioral predictors were *Pr* and *accuracy*. We also included a randomly fluctuating predictor (*~ U*(0,1))) as a control condition. It was denoted as *random*. Each predictor took a turn as the “BOLD signal” (above) onto which all predictors were separately regressed. This round robin procedure allowed us to assess the degree to which each predictor covaried with the others, or was specific. For example, if the first iteration’s BOLD signal was to be *RPE*, the raw RPE trace was convolved with the HRF and white noise (as separate steps). Each of the other confusion predictors (i.e. *value, Pr accuracy, impulse* and *random*) would then be HRF convolved (without noise) and regressed onto the newly anointed RPE-BOLD signal, in turn. That is, each confusion predictor was regressed separately and independently. The regression constants, *t* and *p* for each independent iteration were harvested and retained for later analysis. That is each round robin yielded 5 sets of regression statistics. It is these statistics that are presented in the results. On the next step of the round robin, for example, Pr could be anointed. The raw Pr would then undergo white noise and HRF convolution, and separate regressions by the remaining five predictors (i.e. *value, RPE accuracy, impulse* and *random*). And so on, until all 6 predictors were once anointed as BOLD signals. Once all 6 had their turn that iteration of the simulations terminated, leading to either new random generation of behavioral data or program termination.

For the *separation* analysis we focused on the comparing *value, RPE* over a range of *α* values. Regressions were similarly round robin, but within predictor, e.g. every *RPE* at every *α* took a turn as the BOLD signal, where it was predicted by every other *RPE*, itself included. A *random* condition was included as well.

We took two approaches to the design matrix. In the first, model-based predictors were regressed directly onto the BOLD signal, akin to a simple correlation. This model-only approach serves as a worst-case specificity scenario. The second impulse-and-model design improves specificity by including both an impulse and a model-based regressor, but orthogonalizing the former with respect to the latter.

Computational model and impulse regressors are often correlated or collinear, violating the independence assumptions implicit in the Ordinary Least Squares (OLS) algorithm we used in our SPM procedure. To rectify this we orthogonalized the model-based predictor with respect to the impulse predictor. The orthogonalization procedure is regression-based, wherein the computational model’s predictor is regressed onto the impulse, returning the residuals. These residuals therefore contain only variance present in the computational model. So when this residualized model-based predictor and the impulse are combined into a single design matrix (thus creating our “impulse-and-model” matrix) the impulse regressor captures binomial (i.e. on/off) activity while the other, model-based, regressor picks the computational models contribution. In our results using both impulse and model predictors we present *t*-values from the model predictor, consistent with the model-only results.

This impulse-and-model is an often recommended strategy for doing model-based fMRI. But the model-only design has significant expositional value. First, it is the simplest and most direct route to carrying out a model-based design. This makes it worth examining on its own. Second, its presence serves to highlight the degree of the specificity problem before any corrective action is taken.

In some reward learning analyses stimulus/response and outcome are separated by a short pause, typically 1–4 seconds. While we did not include such a break in the simulated behavioral data we did examined the effect of shifting value and RPE predictors by up to 3 TRs. A delay between the two regressors did increase specificity. It did not do so in a way that qualitatively changed our results. We’d expect even longer delays to have more pronounced effects, but such delays are not common in current designs and would represent a significant experimental temporal opportunity cost when other specificity increasing methods are available (see *Discussion*).

If a researched is concerned with covariates effecting the specificity of there result, it would be typical to included such regressors in the analysis. We don’t do so here because the specificity we’re concerned with isn’t the known co-variates, but instead the many unknown or unconsidered (to the researcher) theoretical alternatives which happen to be weakly collinear with a known (and included in the model) covariate. In such a situations a researcher might mistakenly conclude in favor of the known covariate. This is akin to the inference, upon achieving significance with, say, the “value” covariate, that it is the “correct” model of activity in a given voxel, and so implicitly other models are wrong.

### Defining specificity

As noted in the introduction, the success of impulse-based designs suggests imperfect specificity. So at what point should non-specificity become a concern? We could find no prior work assessing fMRI model specificity. So we pragmatically chose two conservative cut-offs – the 50% and 25% marks. If 50% of the simulations were significant for “false” predictors (i.e. significant when the BOLD signal was not the same as the predictor) we believe these should be considered “indistinguishable”. If five times the standard false positive rate (5%) were falsely significant (i.e., 25%) we posit this should be considered “non-specific”.

## Results

The specificity of model-based fMRI was examined in two analyses. *Confusion* analyses examined the specificity between theoretically distinct (covariate) regressors. The second, *separation*, analyses examined specificity as function of model parameter selection.

### Confusion

Tabulating over all 40 *confusion* analyses, including both modes of simulated learning, the five trial-level predictors were indistinguishable 48% of time and non-specific 52% of time, when using the model-only type design matrix and the *p* < 0.05 cutoff (see *Methods* for details). For the impulse-and-model design overall specificity did improve approximately 15% compared to the model-only design; 34% of the time predictors were indistinguishable and 42% they were non-specific (*p* < 0.05 cutoff).

Overall the impulse-and-model approach does increase specificity. This increase comes at the cost of power. In the model-only design when BOLD signal and predictor match percent significant was ~ 100%. But in impulse-and-model this self-recovery percentage drops to between 15 and 100%, averaging ~80% (compare **A** to **B**, Figures 2 and 3). This reduction in power is despite a uniform decrease in variance between model-only and impulse-and-model distributions (average difference was 1.01*SD, SD_impulse−and−model_* = 2.97, *SD_model−only_* = 3.69) (compare **A** and **B** in Figure 2).

**Figure 2.**
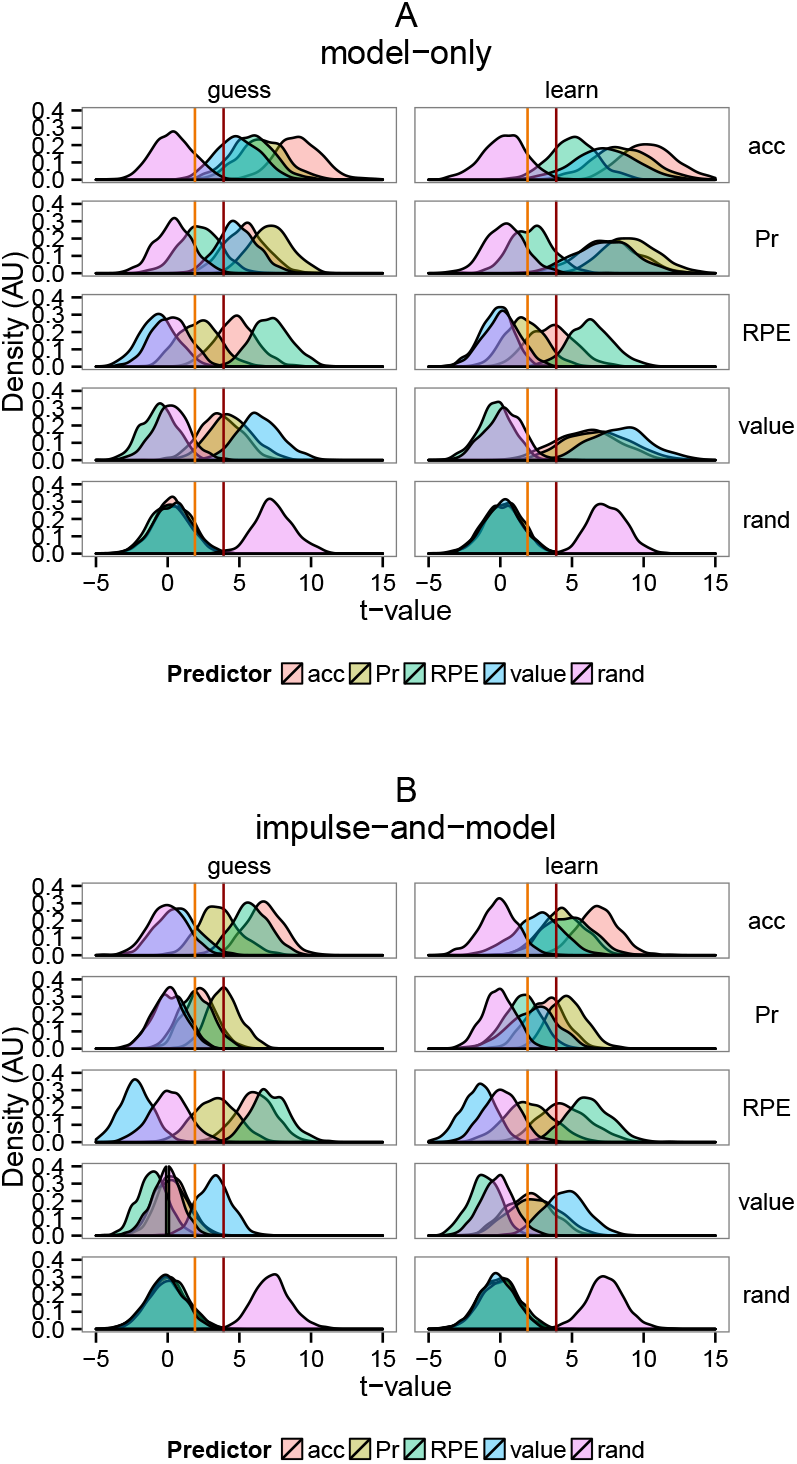
Distribution of *t*-values for all *confusion* analyses, which examined the specificity of model-based fMRI between theoretically distinct trial-level predictors shown in Figure 1**A**. The five predictors were *accuracy* (red), *P_r_* (yellow), *RPE* (green), *value* (blue), and *random* (violet, ‘rand’). In the left column is guessing behavior, on the right is the learning. Each row represents a different true BOLD signal (see labels on the right). That is, each predictor (with noise added) was used to form the simulated BOLD signal that the predictors were fit to. Note that, as expected, there was good correspondence between the actual underlying BOLD signal and the best fitting predictor. Note also the high degree of overlap between the actual predictor and the other predictors across all conditions except *random*. The red and orange vertical lines represent two common statistical cutoffs, *p* < 0.0001 and *p* < 0.05 respectively. For example, in the top row, where accuracy (‘acc’) is the true signal, a large proportion of the distribution for all other predictors other than *random* falls to the right of the p value thresholds, indicating a lack of specificity and a high degree of confusion. The top panel (**A**) represents the model-only approach to GLM regression, while the bottom (**B**) represents the impulse-and-model approach, a commonly recommended procedure for model-based analyses.

Discussing the model-only results in detail, BOLD models based on *accuracy* and *P_r_* were indistinguishable for all other predictors, all that is but the *random* predictor (Figures 2**A** and 3**A**). The *random* was and should have been specific in all cases because it contains no trial-level information, sampled as it was from *U* (0,1), the uniform distribution. It worth noting however that the *random* BOLD model was significant 15–20% of time (see *Discussion*). The *RPE* BOLD model was indistinguishable from *P_r_* and *accuracy* under guessing behavior and indistinguishable and non-specific during learning while *value* was specific (compare left and right columns, Figures 2**A** and 3**A**). When instead *value* acted as the BOLD signal, both *P_r_* and *accuracy* were indistinguishable for both behavior types (Figures 2**A** and 3**A**). Overall, the behavioral model had little effect on specificity (compare columns in Figures 3, see exceptions above).

In the impulse-and-model design, like the model-only design, *accuracy* and *P_r_* BOLD models were non-specific or indistinguishable to all other predictors (Figures 3**B**). *RPE* was indistinguishable from both *accuracy* and *P_r_*. Finally, under the impulse-and-model design only *value* displayed consistent specificity under the guessing behavior (right column, Figure 3**B**). Under learning behavior though specificity was identical to the model-only condition; *P_r_* and *accuracy* were indistinguishable (left column, Figures 2**B** and 3**B**).

**Figure 3.**
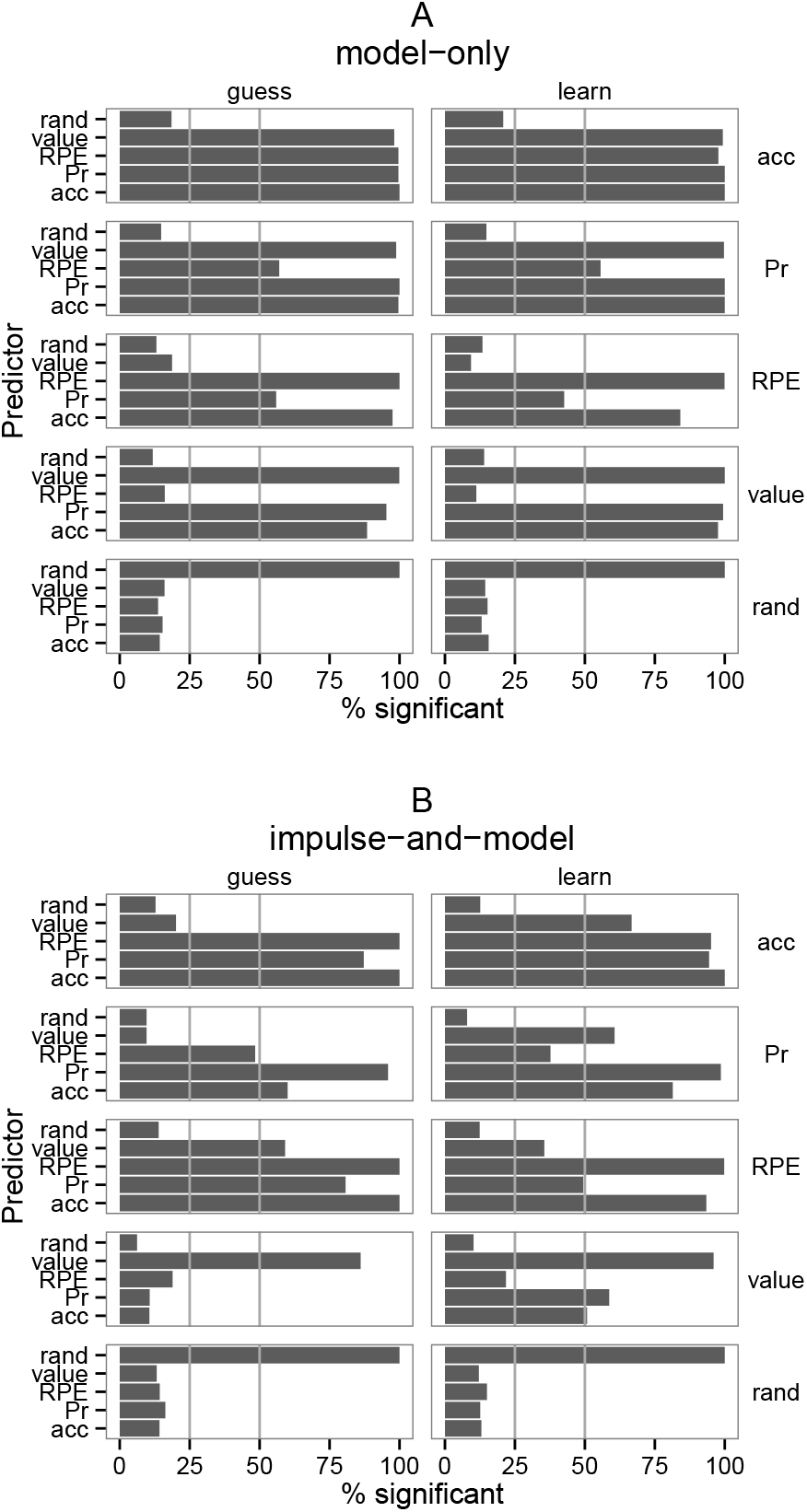
Percent of *confusion* analyses that were significant at the 0.05 level inFigure 2. The two grey lines at 25 and 50% demarcate the cutoffs used to define non-specific (25%) and indistinguishable (50%) results.

### Separation

The *separation* analysis examined specificity between reinforcement learning model learning rate (*α*) parameterizations. No parameter setting was specific. Only the extreme settings of *RPE* were merely non-specific (column 0.1 compared to 0.9, and its complement, Figure 4**A** and **B**). All other *RPE* and *value* predictors were indistinguishable (Figure 4**A** and **B**). The *RPE* values did show an ordering, percent significance increased as predictor and BOLD parameters approached each other. Except for 0.1, *value* showed essentially no such ranking (Figure 4**A** and **B**).

**Figure 4.**
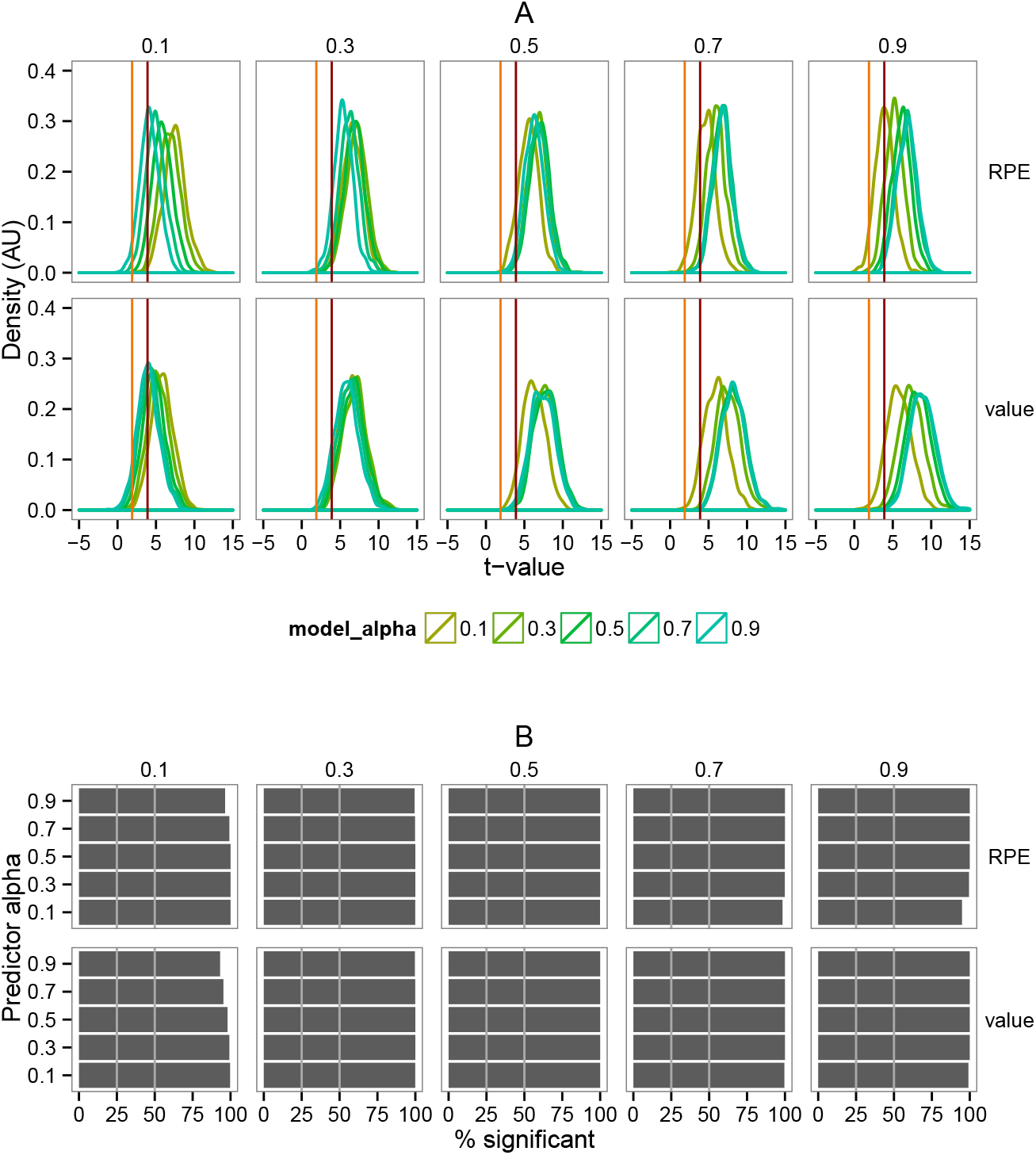
Distribution of *t*-values for both *RPE* and *value* as a function BOLD signals defined using a range of learning rates (see column labels). As in the *confusion* analyses, every alpha value was used in turn to form the underlying BOLD signal, in round-robin fashion. The red and orange vertical lines represent two common statistical cutoffs, *p* < 0.0001 and *p* < 0.05 respectively. **B** Percent of tests from **A** that were significant at 0.05 for each alpha value. The two grey lines at 25 and 50% demarcate the cutoffs we used to define non-specific (25%) and indistinguishable (50%) results. Only guess behavioral models and model-only designs were employed in this analysis.

### Before and after HRF

To separate intrinsic covariance between regressors and HRF induced correlations we measured the Pearsons correlations between 100 randomly selected predictors before and after HRF convolution. In the left column of **A** and **B** in Figure 5 are distribution estimates for raw reward and reinforcement predictors. In the right column are those same predictors after HRF convolution. Following convolution, overlap between distributions increases substantially, while the unique character of the distributions is abolished. For example, compare the bimodal shape *RPE* before convolution to its shape after (Figure 5**A** and **B**). Nor are experiment-level trends in the time courses preserved. Experiments using guess behavior were as non-specific as learning behavior (compare Figure 5**A** and **B**).

**Figure 5.**
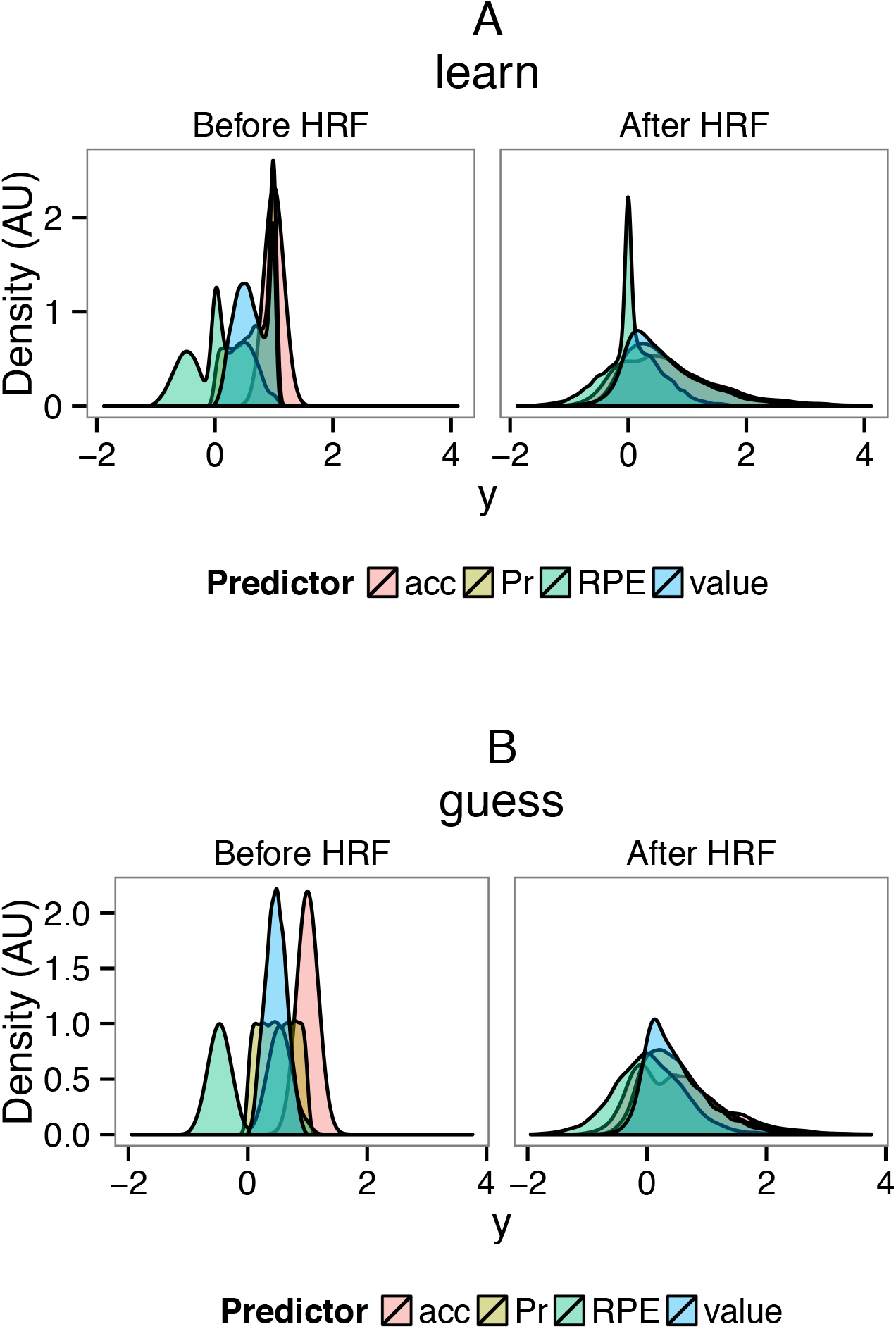
The distribution of 100 randomly selected reinforcement learning time courses, before and after HRF convolution. These are raw data, not distributions of *t* values as in the above figures. The four predictors were *accuracy* (red), *P_r_* (yellow), *RPE* (green), and *value* (blue).

The HRF increased correlations in the majority of pairs, covering a range –0.16 to 0.73 with an average of 0.27 (see Fig 7) The HRF increased correlations among all predictor pairs up until their pre-HRF correlation approached 0.5, after which the correlation began to decline slightly (Fig 6 and Fig 7). When the correlations were broken down into quartiles (Fig 6B), each set of lines had similar slopes (excepting the transition near 0.5) which suggests the HRF has a consistent effect independent of predictor pair. Linear regression analyses supported this conclusion, indicating that *r_before_* could significantly predict *r_diff_*), the difference in correlation before and after, (*F*(1,498) = 713.4, *p* < 2.2*e* − 16) accounting for 0.58 % of the variance. However by including a ‘pair’ dummy predictor (facet labels in Fig 6**A**), a combined pair-*r_before_* model could account for 0.9207 of the variance (*F* (5,494) = 1160, *p* < 2.2*e* − 16) and was a significant improvement over the model using only *r_before_* (*F* (2,494) = 523.5, *p* < 2.2*e* − 16). In total then, the initial correlation between predictors does play a significant role in predicting the change induced by the HRF, however there is a pair (and therefore model) specific component as well (estimated here to be 38% of the total explained variance). A similar analyses of the *separation* analyses data showed a nearly identical pattern (not shown). Follow up analyses of the pair-specific contribution was without significant result.

## Discussion

Using reinforcement learning models as a case study we examined the specificity of model-based fMRI. In the first analysis, dubbed *confusion*, we examined how reliably we could distinguish between related but theoretically distinct predictors. About half the time, the different predictors were indistinguishable (see Figure 2**A** and Figure 3**A**). In the second analysis, dubbed *separation*, we determined that nearly all reinforcement learning model parameter settings were indistinguishable (Figure 4).

### Minimum specificity

In both model-only and model and impulse design matrices 15–20% of guessing behavioral trials were significant (Figure 3**A** and **B**; percents calculated using the *p* < 0.05 threshold). Predictor type had little to no effect on this rate (compare y-axis of row ‘rand’, Figure 3). As the *random* predictor contains no consistent trial-level information we would expect essentially no correlation, and in fact the average correlation between *random* and the other time courses was less than 0.04. Based on this, we conclude that HRF convolution reduces specificity by a minimum of 15%. It should be noted however that 15% is a minimum estimate. By acting as a low-pass filter the HRF may amplify any trial-level covariance (see Figures 6 and 7 for examples and caveats).

**Figure 7.**
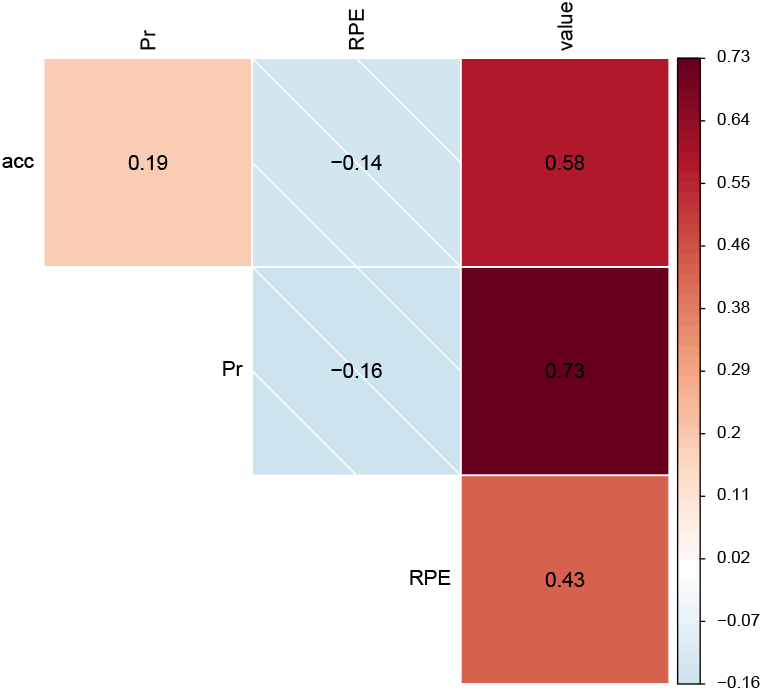
Average difference of predictor correlation matrices before and after HRF convolution, a quantification of overall effect of HRF convolution.

### The impact of the design matrix

Randomization, slow-event related designs, and inter-trial jitter are routinely employed in impulse-based designs to ameliorate correlated regressor issues. The continuous valued and the generative process-derived nature of model-based data limits randomization options. The correlation analysis in Fig 6 suggest that the limited specificity we report is not due to undersampling, or detection power, but is an intrinsic property of HRF convolution implying slow-event related designs would offer little improvement. Likewise, inter-trial jitter works by randomly separating out conditions, allowing for the independent estimation of each condition’s response. However the continuous nature of model-based predictors doesn’t fit well into a condition oriented analysis.

**Figure 6.**
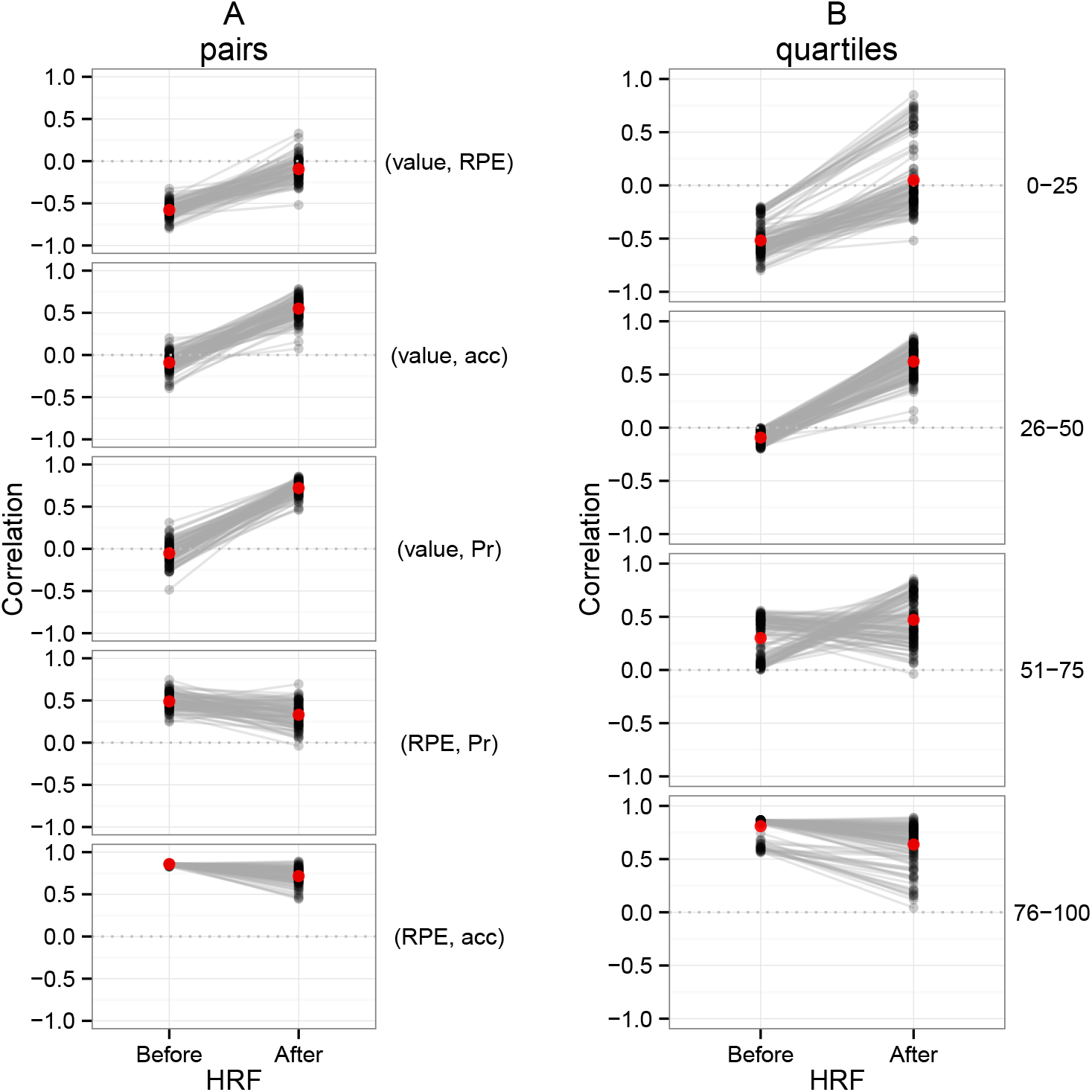
Correlations between each unique pair of predictors (labels in parenthesis) for 100 randomly selected simulations. **B**. The same pairs now binned by quartiles derived from the before HRF convolution set (column labels indicate quartile). The red dot represents the average for that condition.

### Alternatives to OLS

In these simulations it was often the case that the true model was significant more often than the alternatives, even though all were frequently significant (Figure 2). As a result, a model selection or comparison approach to model-based fMRI should be fruitful. Model selection is the process of finding a *family* of models that best predict a given dataset (Rao et al., 2001). This can be a simple as comparing the explained variance (i.e. *R*^2^) between model options. More sophisticated techniques though will attempt to balance parsimony with increasing fit (i.e., solving the bias versus variance dilemma (Geman et al., 1992)). Examples include Stepwise, LASSO, or Ridge (Tikhonov) regression. Ridge regression is of particular interest as it is capable of selecting among covariate or collinear predictors, as was the case for our reinforcement and reward-related predictors. As an alternative to algorithmic selection procedures, and the sometimes stronger assumptions they entail, candidate models could be directly compared using standard model comparison metrics, such as Akaike Information Criterion (AIC) weights (Wagenmakers and Farrell, 2004). In doing to model comparisons however, it is crucial that one selects suitable and viable alternate models, specifically avoiding “stacking the deck” in favor of a preferred option.

### Problems for parameters

Given that reinforcement learning parameters are typically set based on behavioral data, one might first assume it is not important that the GLM procedure cannot distinguish between parameters. However, further reflection suggests the lack of parameter sensitivity is important for three reasons. First, given a significant result it would normally be tempting to conclude that, “our model was a significant predictor of the BOLD changes under parameters *{α,β,θ}* therefore neural activity may reflects these models and parameters”. But given our results it quite possible a very different set of parameters, *{α′,β′,θ′}*’, would also be significant. Without specificity all a finding of significance can guarantee is that, “Some (unspecified) set of parameters near our parameters *{α,β,θ}* were significant predictors of BOLD activity”, which seems a deeply unsatisfying *best case* conclusion. Second, parameter changes can considerably alter model behavior. For example, in Figure 1**D** the choice of the learning rate parameter *α* affected both qualitative (compare 0.1 with 0.9) and quantitative (compare 0.1 with 0.3) behavior. Thirdly, model-based designs are often used in the clinical literature to examine neural correlate differences between patient and control populations (Castro-Rodrigues and Oliveira-Maia, 2013; Deserno et al., 2013). Without specificity, such comparisons are meaningless.

Consider again the *separation* data (Figure 4); it would be difficult for any method to separate the overlapped distributions observed, unless a very large number of samples were available or only extreme parameters settings are considered. If true beyond our case-study (as may be the case, see Fig 6), reliable parameter separation in model-based fMRI may prove quite difficult. Computational models are useful, in part, because they make quantitative predictions, predictions which depend on parameter settings. If you can’t distinguish between parameters, you can’t distinguish between quantitative predictions.

### Limitations and generalizations

The kind of specificity we study here is most relevant when two or more theoretical models are compared, and when predictions depend on parameter choices. This does not mean that the specificity issues are limited to computational modeling. Any parametric design, for example one derived from reward value, reaction time, or subjective scoring, like happiness or emotional salience, may also have it’s trial-level specificity diminished by convolution with the HRF. In that sense, our case study of reinforcement learning may have much broader potential implications. A reduction in specificity may happen whenever the predictor changes faster then the BOLD response, which acts akin to a low-pass filter.

While low-pass filtering by the HRF is always possible, specificity will matter less when when models or parametric regressors are being used as stand-ins for latent psychological states. Specificity doesn’t matter in cases where parametric responses are estimated from *repeated measurements* of fixed conditions, such the recent retinotopy-based image reconstruction work ?. That is, the specificity problem we begin to demonstrate is not present when the goal is estimate the expectation of some consistent response function, but is present when we the aim is to account for as much trial-level variability as possible. Finally, if several models were compared to the BOLD response and their response were averaged, this would implicitly lessen the importance of any one model’s specificity.

### Conclusions

To study the relation between significance and specificity in model-based fMRI we made a case study of reinforcement learning. Being a case study, and an idealized one at that, the generality of these results to other models and past empirical work is unproven. But what we have found urges caution. The prototyped shape and long temporal evolution of the haemodynamic response function can induce strong positive correlations between predictors. In our worst case, the HRF more than doubled the correlation between predictors, about halving specificity. We urge researchers to consider carefully the interpretation of their significant model-based results and perhaps to move from a null hypothesis testing framework to model comparison framework.

## Disclosure/Conflict-of-Interest Statement

The authors declare that the research was conducted in the absence of any commercial or financial relationships that could be construed as a potential conflict of interest.

## Author Contributions

EJP designed and conducted the study, and wrote the paper. CAS helped design the study and wrote the paper.

## Acknowledgement

We thank Bradley Voytek and Howard Landman for their useful feedback on an earlier version of the manuscript. This work was carried out under National Institutes of Health R01MH079182.

